# Dynamic blinking in the head of hardyhead silverside fish

**DOI:** 10.1101/2021.02.03.429660

**Authors:** Masakazu Iwasaka

## Abstract

Dynamic light reflection can serve a similar purpose to tools such as digital line processing devices. It is interesting, therefore, that evidence of dynamic light reflection can also be found in the animal kingdom and that there may be alternative ways of actuating light control. This study discovered that several features contained in the heads of hardyhead silverside fish, particularly around the edges of the iris, caused blinking using environmentally scattered light. Analyzing the blinking using recorded video of the fish iris revealed that circular cells existing in the iris changed their light intensity at 2 Hz. These 5–10-μm-diameter cells are normally blue. However, it is found that a distinct light intensity changed in 0.04 seconds, and additional green and yellow colors overlapped with the blue. It is hoped that utilizing the mechanism that controls the rapid changes in light intensity using only environmental lighting can reduce electrical power usage in display devices.

## Introduction

Light reflection is one of the most interesting phenomena in the fields of optics and photonics and has also been widely reported on in the fields of physics (1), geoscience (2–5), optical engineering (6–10), biophysics (11), and medicine (12–17). By detecting the scattered light at the periphery of incidental light, darkfield illumination has allowed for light particles to be optically analyzed on a nanometer scale (1). Measurements of light reflection have also been applied to larger objects such as our entire planet (2–5). The data collected from light absorption and reflectance by the land and atmosphere are very useful in sensing the environmental condition.

Detailed investigations of light reflection from objects similar in size to the human body can be facilitated by improvements in computer science (6–9). Computer simulation techniques for modeling light reflection by various model objects have become some of the most essential tools in today’s information society. One of these advanced semiconducting technologies, the digital micromirror device, involves applying light reflection to a smooth aluminum mirror, which can now be attached to the human body (10).

Darkfield illumination, a technique developed in the previous century, can now be used at the microscopic level using a contemporary imaging technique (11). Most medical research concerning light reflection was conducted more than 50 years ago (1, 12–17). However, the analysis of light absorption and reflectance in living tissues near the body surface, such as the skin and peripheral blood vessels, is now an essential part of medical diagnoses (12–14). Additionally, light reflected by the pupil is linked to information processing in the brain (15–17). These studies suggest that an optical property, such as the light reflectance of living tissue, is strongly correlated with signal indication and processing in living creatures.

The skin and scales of fish can be used to inspect unrevealed optical properties of the living body exposed to external light because most fish have reflecting platelets in the skin (18). Our previous studies focused on several types of light-reflecting behaviors in the reflective guanine platelets found in fish skin (19–22). In recent years, I studied fish living in the sub-tropical regions of Japan to find any instances of dynamic and tunable light reflection phenomena. In this study, I report on how the head of a particular species of fish, the hardyhead silverside (*Atherinomorus lacunosus*), is able to dynamically reflect light.

## Materials and Methods

### Specimen

Hardyhead silverside fish, *Atherinomorus lacunosus*, (five specimens) were obtained in Okinawa, Japan. After transport to Hiroshima, the fish were maintained in an aerated seawater aquarium with a water temperature thermostatically controlled at 26–29 °C. The five specimens were used in experiments ‘conducted in December 2020 in accordance with the ethics policies of Hiroshima University Animal Care and Use Committee (approval number: F19-2, Hiroshima University).

### Observation of fish in aquarium

The fish in the aquarium were recorded by a lens (***, Nikon, Tokyo, Japan), while the microscopic images of the heads of the anesthetized fish were taken using a high-resolution Navitar 2.0 × 1-51473 (Navitar, Rochester, USA) microscopic lens. Both lenses were connected to a CMOS camera (HOZAN L-835, HOZAN, Osaka, Japan) by a C-mount adapter when recording the fish. The white balance and exposure times were controlled manually and a white LA-HDF158A LED light (Hayashi Repic Co. Ltd., Tokyo, Japan) was used as a light source.

The fish were anesthetized by exposing them to 0.1% 2-phenoxy-ethanol for 10–30 s at approximately 15 °C, which was the temperature of the water in our experimental room. The anesthetized fish were then placed inside a plastic aquarium containing aerated seawater. After a short period of observation, the fish were then returned to the main aquarium.

### Light intensity dynamics measurements on a computer LCD screen during playback of recorded video

Measurements on a computer display were collected using a fiberoptic system. First, the video playback was paused at the place where the measured point appeared. One of the ends of the optical fiber was then placed close to the measured point and fixed onto the LCD screen. The light intensity data were collected by a CCD spectrophotometer (USP-2000, Unisoku Co. Ltd. Osaka, Japan) and the video playback resumed a few seconds after the data collection began.

## Results and Discussion

Figure 1 shows an image recorded from the video of the blinking eye of a hardyhead silverside fish swimming in the aquarium. All the investigated specimens (four samples) had a common part that demonstrated blinking, which appeared intermittently in seven parts of the head (I–V, VIII, and IX in Figure 1A–C) and in two parts of the body (VI and VII in Figure 1A–C). The blinking occurred most strongly and frequently in Parts I, II, and III. Parts I and II were located at the iris edges and Part III was located in the center of the skull at the midpoint of the eyes. In Part IX, it looked as though the small circles were blinking. The angles of incidental light when hitting the fish’s body, the aquarium, and the water surface are illustrated in Figure 1D. Because the fish were swimming in an aerated, thermostatically controlled environment, their bodies were always in motion, even when floating statically.

**Figure 1.**
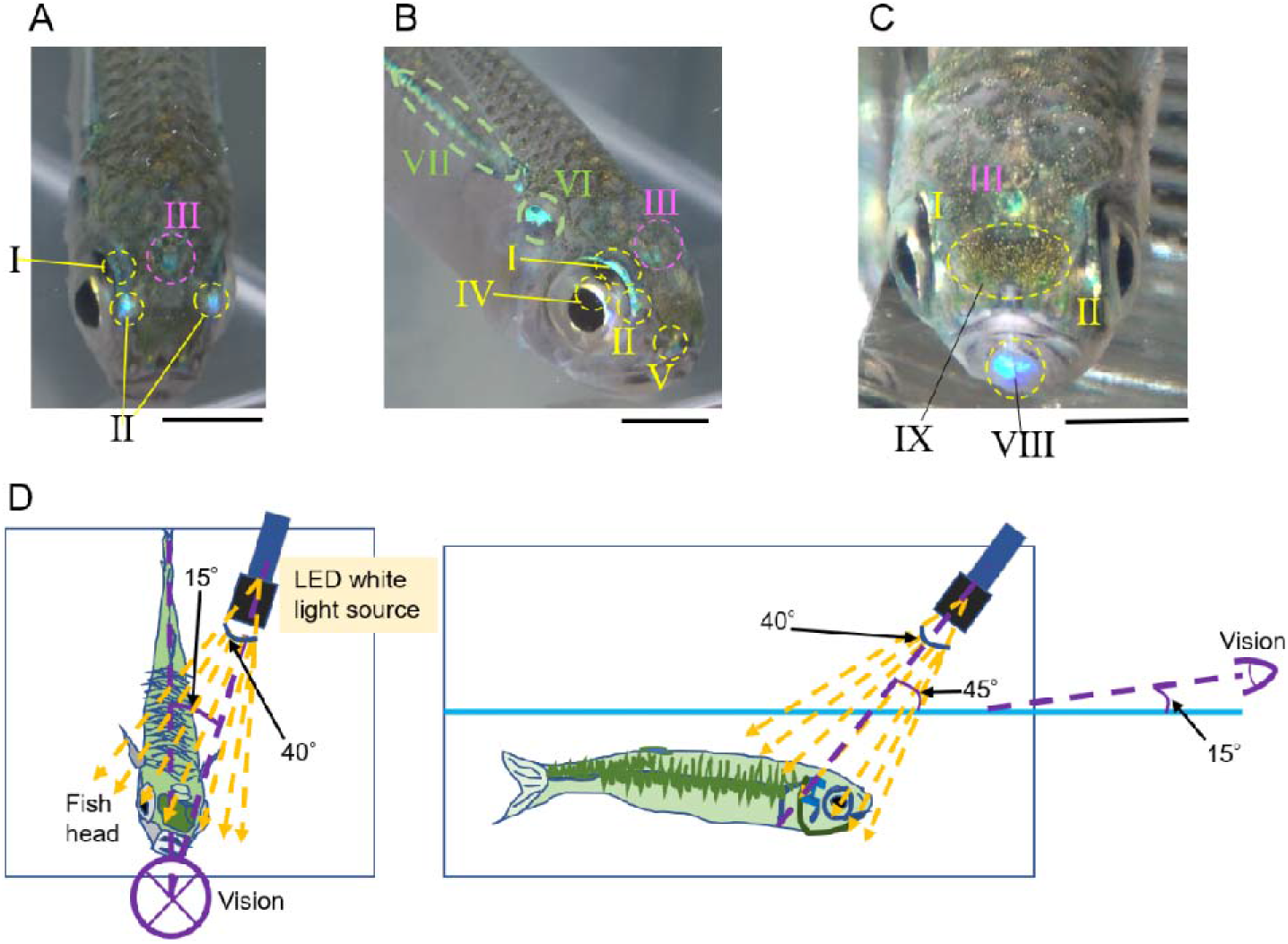
Dynamic blinking parts in body (mostly in head) of hardyhead silverside fish. The blinking seems to be the consequence of the light reflecting by reflectors in each of tissues. (A)-(C), Representative parts (I ~ IX)) showing remarkable blinking. (D), Roughly sketched condition of the exposure of specimen to incident light. The light beam from the light guide expanded and was provided to the body surface with a dispersion of angles up to 40°. Scale Bar is 10 mm.

We can theorize that the blinking on the body side, Parts VI and VII, correlates with the inclination of the body. It was more difficult, however, to distinguish between the effects of body motion and possible actuation in local tissues. Regarding Part VIII, it was initially thought that the blue blinking shown in Figure 1C was caused by changes in light reflection changes caused by the movement of the computer mouse. However, the video images did not corroborate this theory.

Figure 2 shows 18 randomly captured images of the eye of a hardyhead silverside fish swimming in an aquarium (Supporting Movie A). All of the individual images have slightly different inclinations. Remarkably, an increase in light intensity occurred at the frontal edge of the iris (Part II), but the timing did not seem to correlate with the eye’s inclination. The blinking observed in Parts I and II were both detectable by the naked eye.

**Figure 2.**
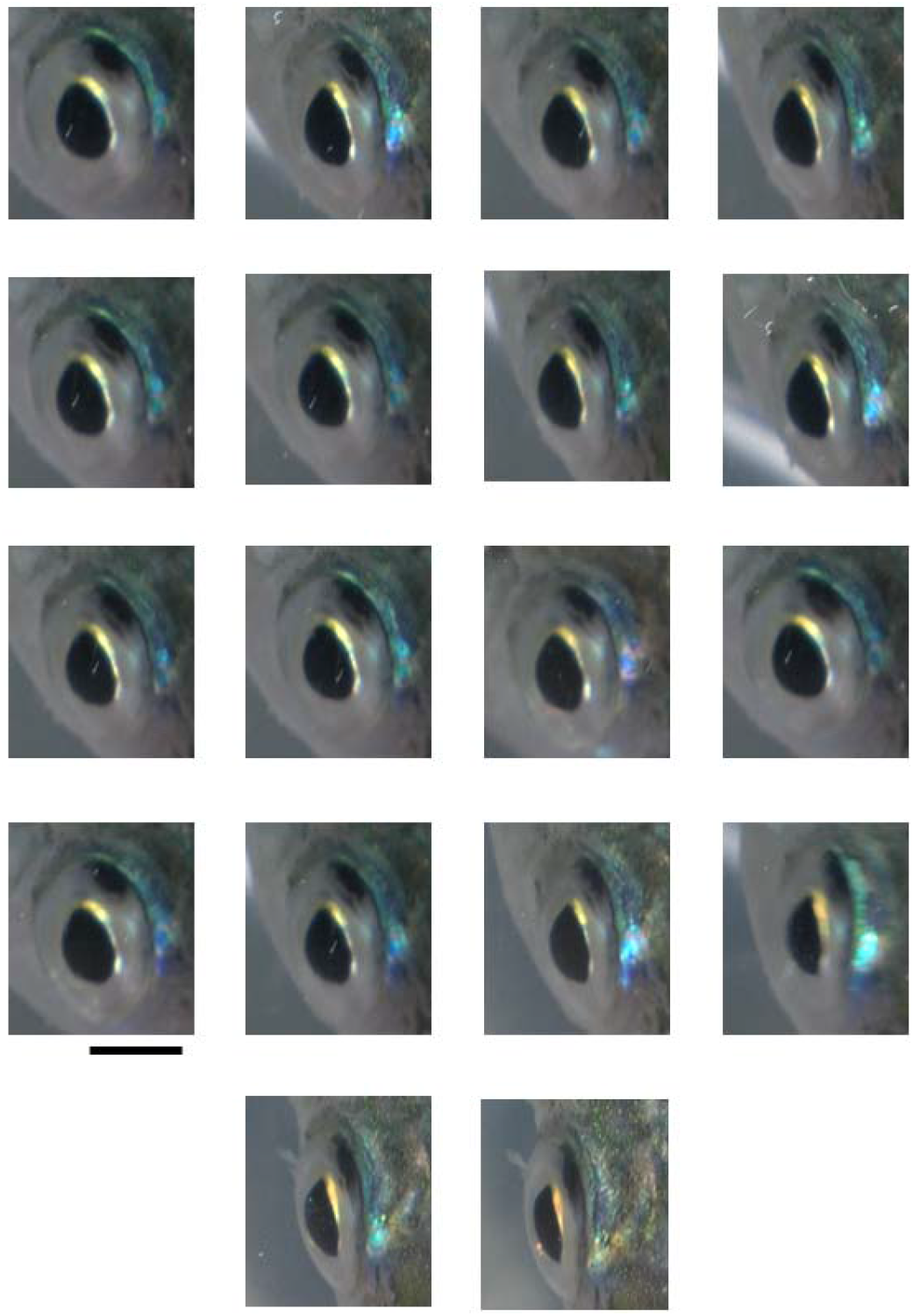
Dynamic blinking parts in edge of iris. Eighteen images were captured from the movie of statically swimming hardyhead silverside fish (Supporting movie-1). Scale Bar is 5 mm.

Next, to investigate whether the blinking at the edge of the iris occurred independently in the local part of the iris, microscopic observation of the edge of the iris (Part I) was conducted on an anesthetized fish. Figure 3 shows two image sequences obtained from the recorded video (Supporting Movies B and C), with the magnification being the only difference between the two sequences. The videos were recorded both while the fish were anesthetized and when they were quietly settled in the main aquarium.

**Figure 3.**
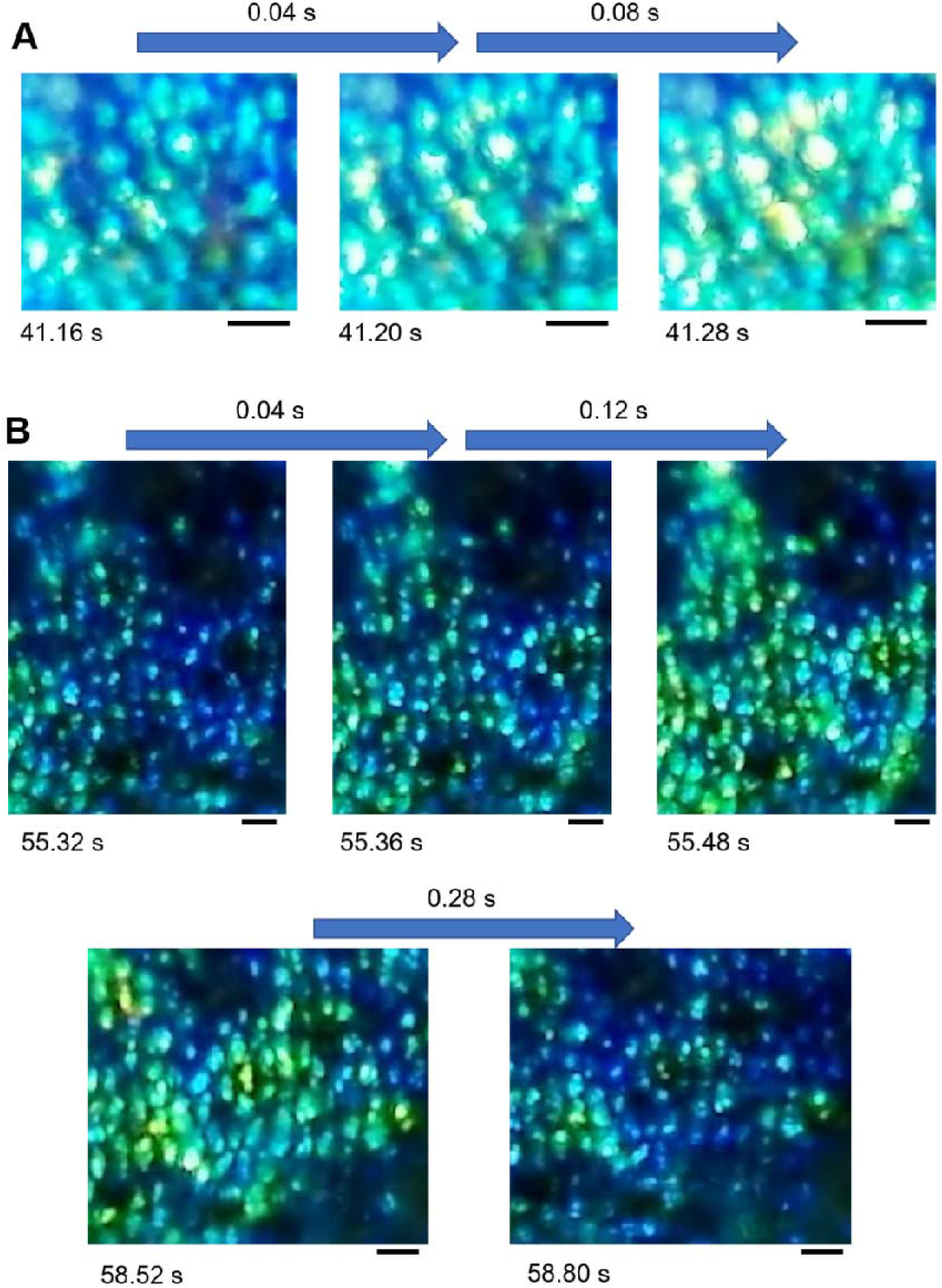
Microscopic observation of blinking in edge of iris (part I shown in Fig. 1 A-C) of anesthetized hardyhead silverside fish. (A) First example showing a distinct light intensity increase in 0.12 s. Time indication shows the playback time in Supporting movie-2. (B) Second example showing increase and decrease of light intensity. Time indication shows the playback time in Supporting movie-3. Scale Bar is 10 μm.

Aggregated circular cells were found on the edge of the iris. The first example (Figure 3A) shows a rapid increase in light intensity after 0.12 s with pale-green circular cells visible on the blue base (left image of Fig. 3A). Among them, approximately eight cells changed their color to yellow (middle image) after 0.04 s, and the color intensity increased after 0.08 s (right image). Figure 3B shows a similar phenomenon, but with less expansion.

The image sequence shows a cycle of increasing and decreasing light intensity over a period of 0.44 s. Initially, most of the circular cells were blue but they turned green or yellow after approximately half a second. These results indicate that the condensed circular cells in the iris were blinking but did not change the inclination of the iris. As shown in Figure 3 and Supporting Movies 2 and 3, the blinking was continuous for one minute, and the frequency of the blinking was approximately 2 Hz. This was confirmed by analyzing the blinking of the iris during playback (Supporting Movie 2), as shown in Figure 4. The fiberoptic measurements on a local area in LCD projecting the video provided time courses of light intensity, as shown in Figures 4B and 4C.

**Figure 4.**
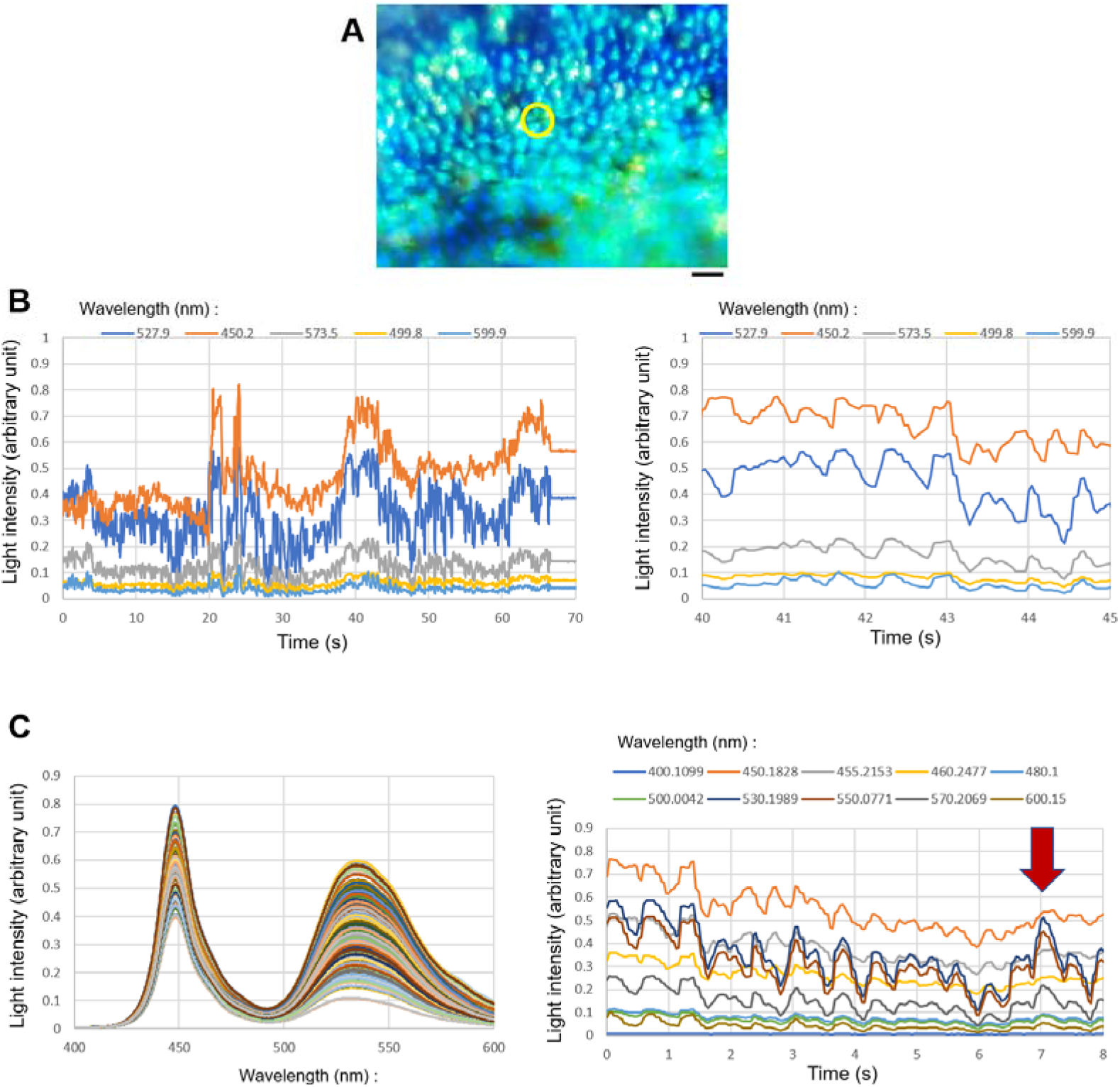
Analysis of frequency of blinking in iris of hardyhead silverside fish by measuring light from a computer LCD screen during playback of recorded video. (A) Measured area (marked by yellow circle) in the video (Supporting movie-2). Scale bar is 20 μm. (B) Time course of light intensity at five wavelengths of blinking area. Right panel shows the expansion of the time course at 40 - 45 s. (C) Measurement of spectrum for eight seconds during the playback of the same video. Right panel shows time course of light at selected ten wavelengths.

Figure 4B shows that the area covering four circular cells exhibited blinking for more than 60 seconds. An expansion of the time course (right-hand panel) indicates that the blinking occurred at a frequency of 2 Hz.

In Figure 4C, the light from the measured area was found to be in the range of 400 nm to 600 nm for approximately 8 seconds, then the data was transferred to the time course style. Among the 10 selected wavelengths, the time courses at 3 wavelengths between 530 nm and 570 nm (dark blue, brown, and intense gray lines) exhibited remarkable oscillations with the same frequency at approximately 2 Hz. The time course at 450 nm (orange line) also shows an oscillation at 2 Hz, although the oscillation pattern was different from those between 530 nm and 570 nm. As shown in the right-hand panel of Figure 4C, the light intensity at the three wavelengths between 530 nm and 570 nm increased dramatically after 0.7 s, while the change at 450 nm was comparatively small.

The blinking circular cells appear to be a type of chromatophore [23], a group of cells that control light absorption and reflection onto the bodies of animals such as fish and amphibians. It is possible that the circular cells, which are less than 10 μm in diameter, are a type of chromatophore called an iridophore. This is because iridophores often contain reflective platelets inside the cell. The circular cells observed in our study clearly changed their light intensity, probably by using reflective particles within the cell.

The light intensity on the eye of hardyhead silverside fish changed after 0.04 s, as shown in Figure 3. The speed of the change was faster than the iridophores found in the bodies of flashing tilefish, which changed color in half a second [24]. It was reported that the color changes in the iridophores found in flashing tilefish were controlled by the nervous system. Whether or not the nervous system controls the blinking in the heads of hardyhead silverside fish will need to be investigated in a future study.

## Conclusion

Hardyhead silverside fish had blinking parts on head. The blinking was obvious when swimming in an aquarium. The most distinctive blinking parts were located on the edges of the iris. By microscopically observing the irises of anesthetized fish, we found that the circular cells on the iris were causing the continuous blinking. A computer LCD screen used during playback of the recorded video confirmed that the blinking occurred at a frequency of 2 Hz.

## Authors’ contributions

M.I. performed the experimental design, experiments, measurements and analyses. All part of manuscript and illustrations were prepared by M. I.

## Acknowledgments

Author appreciates Y. Uehara for providing the specimen of fish.

## Supporting Information

**Supporting Movie A: Hardyhead silverside fish swimming in an aquarium**

(Real-time movie).

This movie provided the images for Figures 1 and 2. Detailed conditions about the observation can be found in the main body. The original movie was in AVI format, but the file was converted to MPEG-4.

**Supporting Movies B and C: Real-time movies of the blinking parts on the edge of the iris in hardyhead silverside fish.**

These movies provided the images for Figure 3. Detailed conditions of the observation can be found in the main body. The only difference between the two movies is the magnification. The videos were recorded both while the fish were anesthetized and when they were quietly settled in the main aquarium. The original movies were in AVI format, but the files were converted to MPEG-4.

